# Antarctic marine microplastics reveals environmental persistence and rapid evolution of *Candida auris*

**DOI:** 10.64898/2026.03.13.711634

**Authors:** Norman van Rhijn, Emma Gan, Pilvi Hepo-oja, Xinyi Wang, Jia Li, Seána Duggan, David Firer, Laila Alsharqi, Hugh Gifford, Jacob L. Steenwyk, Amélie P. Brackin, Alireza Abdolrasouli, Andrew M. Borman, Christina A. Cuomo, Matthew C. Fisher, Darius Armstrong-James, Rhys A. Farrer, Jane Usher, Johanna Rhodes

## Abstract

*Candida* (*Candidozyma*) *auris* is a critical priority fungal pathogen that emerged two decades ago near simultaneously on multiple continents. Since emergence, *C. auris* resistance to all four classes of antifungal drugs has been described, including pan-drug resistant isolates, sometimes evolving within patients. Here, we confirm the first isolation of *C. auris* from Antarctica and show cold-adapted phenotypes and an affinity for binding to nylon. We also provide evidence to suggest mutator phenotypes contribute to the rapid evolution of *C. auris* and are responsible for the emergence of multiple, distinct genetic clades worldwide. Isolates in clades I, III and IV with a mutator phenotype displayed elevated mutation rates compared to non-*auris Candida* species. This phenotype had a complex genetic basis and was associated with drug resistance mutations. We postulate that the mutator phenotype has a significant effect on evolutionary potential and is responsible for the emergence and rapid spread of drug-resistance *C. auris* and novel genetic clades.

## Main

Since the first description of *Candida* (*Candidozyma*) *auris* in Japan in 2009 ^1^, multiple genetically distinct lineages, termed Clades I-VI, have been identified worldwide and reported in over 60 countries ^2,3^. Of these, Clades I, III and IV routinely cause nosocomial outbreaks, with high rates of patient-to-patient transmission and rapid acquisition of antifungal drug resistance ^4–7^. The strong population structure, dominated by deeply diverging genetic clades with high intra-clade clonality, is reinforced by apparent low levels of recombination between clades ^8^. It has been hypothesised that the ancestral population evolved virulence and diversified ^9^, perhaps through sympatric evolutionary processes, before independently emerging. Therefore, an ancestral *C. auris* population may have been exposed to elevated temperatures and other selective pressures, possibly because of climate change ^10,11^, resulting in adaptation facilitating human infection through the accumulation of mutations.

Despite extensive surveillance, the origin and transmission routes of *C. auris* remain unclear. Several studies have recovered *C. auris* from marine environments¹D, and its close relative *Candida haemulonii* also appears to have a marine origin¹¹, although environmental infection rates remain low^12^. *C. auris* demonstrates enhanced thermotolerance and halotolerance compared to phylogenetically related species ^11^, further supporting the hypothesis that this fungal pathogen previously existed as an environmental generalist in areas of high salinity and impacted by climate warming ^10^. Given the propensity for *C. auris* to survive on plastic surfaces for two to four weeks ^13,14^, it is possible that marine plastic pollution could provide a means of worldwide dissemination. Given the ability of *C. auris* to persist on plastic surfaces, marine plastic debris may provide a durable substrate for long-distance dispersal. We therefore sought to test the hypothesis that plastic-associated marine environments could act as reservoirs facilitating global dissemination of *C. auris*.

### *Candida auris* is present in Antarctic marine environments

Although *C. auris* has been sampled across most continents and diverse environmental settings, only seven studies to date have successfully recovered the pathogen from non-clinical environments^15–20^. Motivated by the proposed marine origin of *C. auris* and its notable ability to persist on abiotic surfaces, we collected microplastic debris (ranging from 5 mm to 63 µm in size) using surface sampling approaches to minimise contamination from water or human interference. Samples were taken from six Antarctic marine locations, each separated by an average of 150 km to minimise sampling bias from a single source (Figure 1a).

**Figure 1:**
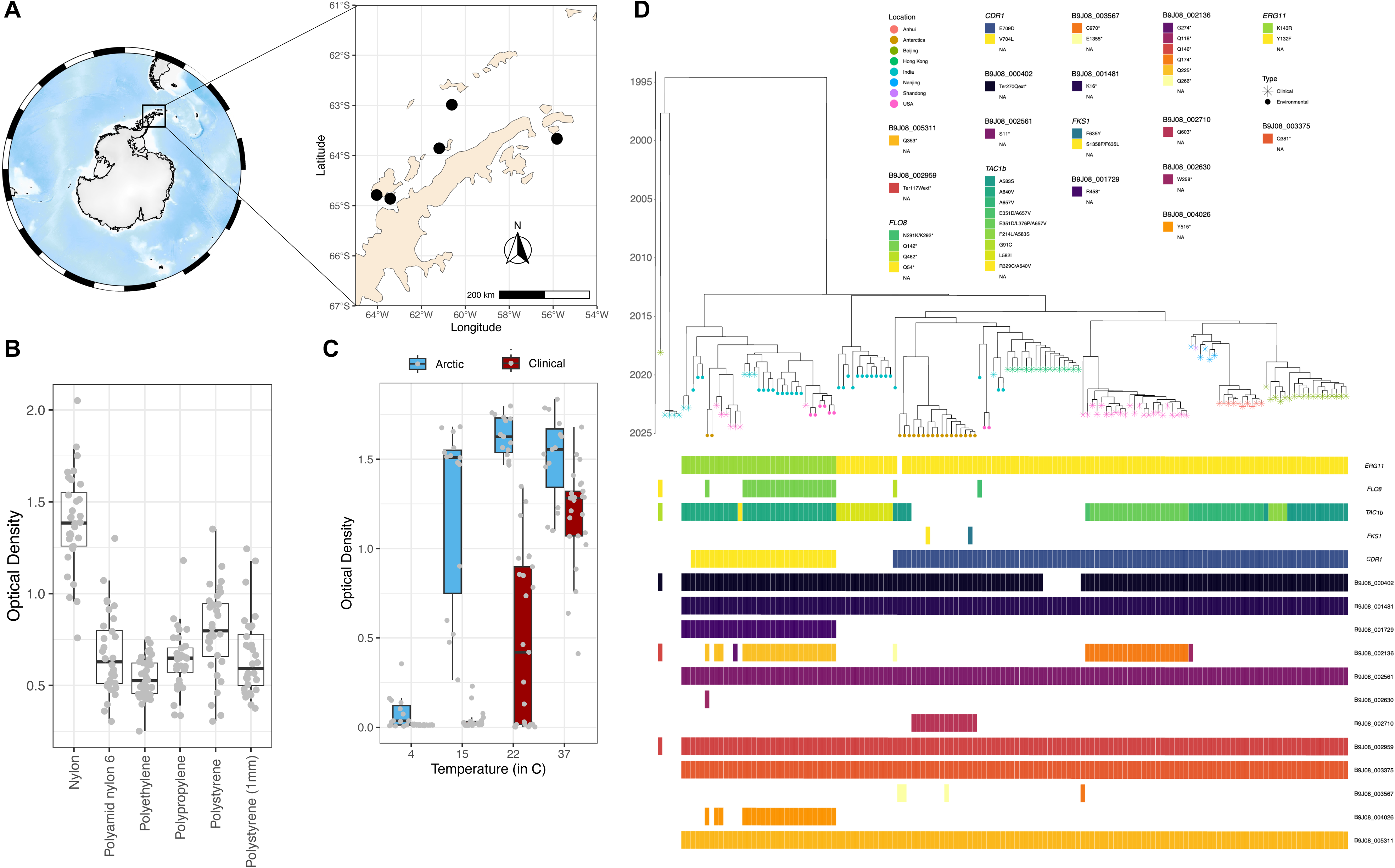
*C. auris* in Antarctic marine environments. **a)** Map of Antarctica and inset with sampling locations (black dots) **b)** *C. auris* biofilm production on various plastic substrates demonstrates preference for nylon **c)** Growth of Antarctic and Clade I clinical *C. auris* isolates shows adaptation of Antarctic isolates to cold temperatures **d)** Time-scaled phylogeny of Antarctic isolates plus publicly available whole genome sequenced environmental and clinical isolates with nonsense and missense mutations in key AMR genes. Tip shape denotes source (clinical or environmental), tip colour denotes sampling location. Nonsense mutations in genes, encoding truncated proteins, are provided in the heatmap.

We recovered nineteen *C. auris* isolates across these six Antarctic marine locations. Phenotypic characterisation revealed that all isolates rapidly formed robust biofilms on nylon (OD=1.392, *P*≤0.0001) and polystyrene (OD=0.798, *P*=0.0002), with markedly lower biofilm formation on other substrates (Figure 1b). These Antarctic isolates also exhibited enhanced growth at lower temperatures than clinical *C. auris* isolates (*n* = 25, *P*≤0.0001 at 22°C, *P*≤0.0001 at 15°C), suggesting they have adapted to cold marine environments (Figure 1c).

Whole genome sequencing and phylogenetic analysis placed all 19 Antarctic isolates within Clade I based on phylogenetic placement (average 133x coverage with 99% of the B8441 v3 reference genome covered, Supplementary Table 1). Pairwise comparison of highlZlconfidence SNPs revealed moderate genetic diversity among the Antarctic isolates (72 SNPs [SD=47]) (Supplementary Figure 1). All Antarctic isolates contained canonical Clade I *ERG11* resistance-conferring mutations Y132F (*n* = 17) or K143R (*n* = 2) (Supplementary Table 1; Figure 1d), which resulted in azole resistance (Supplementary Figure 2). Two isolates (11%; C.aurANT_18 and C.aurANT_6) additionally carried *FKS1* mutations (F635L and F635Y), both of which showed echinocandin resistance (Supplemental Figure 2). Two missense mutations (E709D; *n* = 17, and V704L; *n* = 2) were observed in non-overlapping isolates in *CDR1*, and two missense mutations (A583S; *n* = 3 and A640V; *n* = 2) were observed in *TAC1b* in five different isolates. One isolate, C.aurANT_17 also contained a nonsense mutation (Q142*), encoding a premature stop codon, in *FLO8*, which has been implicated in amphotericin B resistance and biofilm formation (Figure 1d).

Phylogenetic analysis incorporating environmental and clinical reference isolates indicated two independent introductions of *C. auris* into Antarctica (Figure 1d). A single introduction ([2016,2019], 95% HPD; Supplementary Figure 2) of two Antarctic isolates with the *ERG11* K143R mutation were members of one cluster with isolates from India in a basal placement. A second, earlier introduction ([2014,2016], 95% HPD, Supplementary Figure 3) of 17 Antarctic isolates with the *ERG11* Y132F mutation were placed in a cluster with basal Indian and Hong Kong isolates.

We detected missense mutations in three mismatch repair (MMR) pathway genes, *RAD18* (Q145R), *MRE11* (L391V), and *RAD23* (S99C), in all of the Antarctic isolates (*n* = 19). This prompted us to investigate the broader prevalence and functional implications of MMR associated mutations across the global *C. auris* population.

### Clade-specific mutational profiles in MMR pathway genes

Mutations that impair the DNA mismatch repair (MMR) pathway can increase mutation rates across diverse organisms, including bacteria, fungi, and mammals ^21^. In pathogenic fungi, MMR deficient strains frequently display a hypermutator phenotype that accelerate adaptive evolution, including antifungal drug resistance ^22–25^. However, elevated mutation rates also lead to the accumulation of deleterious mutations, resulting in a fitness cost in stable environments. We recently identified an elevated mutation rate (5-fold) resulting in a mutator phenotype with no associated fitness cost in *Aspergillus fumigatus*, found exclusively in isolates harbouring mutations conferring azole resistance ^26^. The underlying genetic basis involved multiple interacting mutations across core MMR genes *msh6*, *msh2* and *pms1*.

To determine how widespread MMR-associated pathway mutations are the wider *C. auris* population, we assessed gene synteny, followed by annotation of MMR genes between the reference genomes of Clades I-V by detecting syntenic gene blocks (Figure 2a) ^28^. The genomes of the five clades are highly syntenic, as described in previous studies ^29^. We also observed evidence of large chromosomal rearrangements between Clade I, II and V (Figure 2a).We recently compiled a dataset of over 12,000 publicly available WGS consisting of Clades I-VI, which we used to assessed clade-specific mutations in MMR genes ^27^. In total, 505 missense mutations were identified in MMR pathway genes relative to the B8441 v3 reference genome (484,284 total mutations identified over full dataset, 0.1%). 31 insertion or deletion frameshift (in *RAD* genes, *MSH2*, *MSH3*, *MLH3*) and two nonsense mutations (in *RAD9* and *RAD26*) were also detected. Missense mutations were present in *MSH*, *RAD* and *Mre11* genes. This pattern suggests that most variants modulate, rather than fully disrupt MMR function.

**Figure 2:**
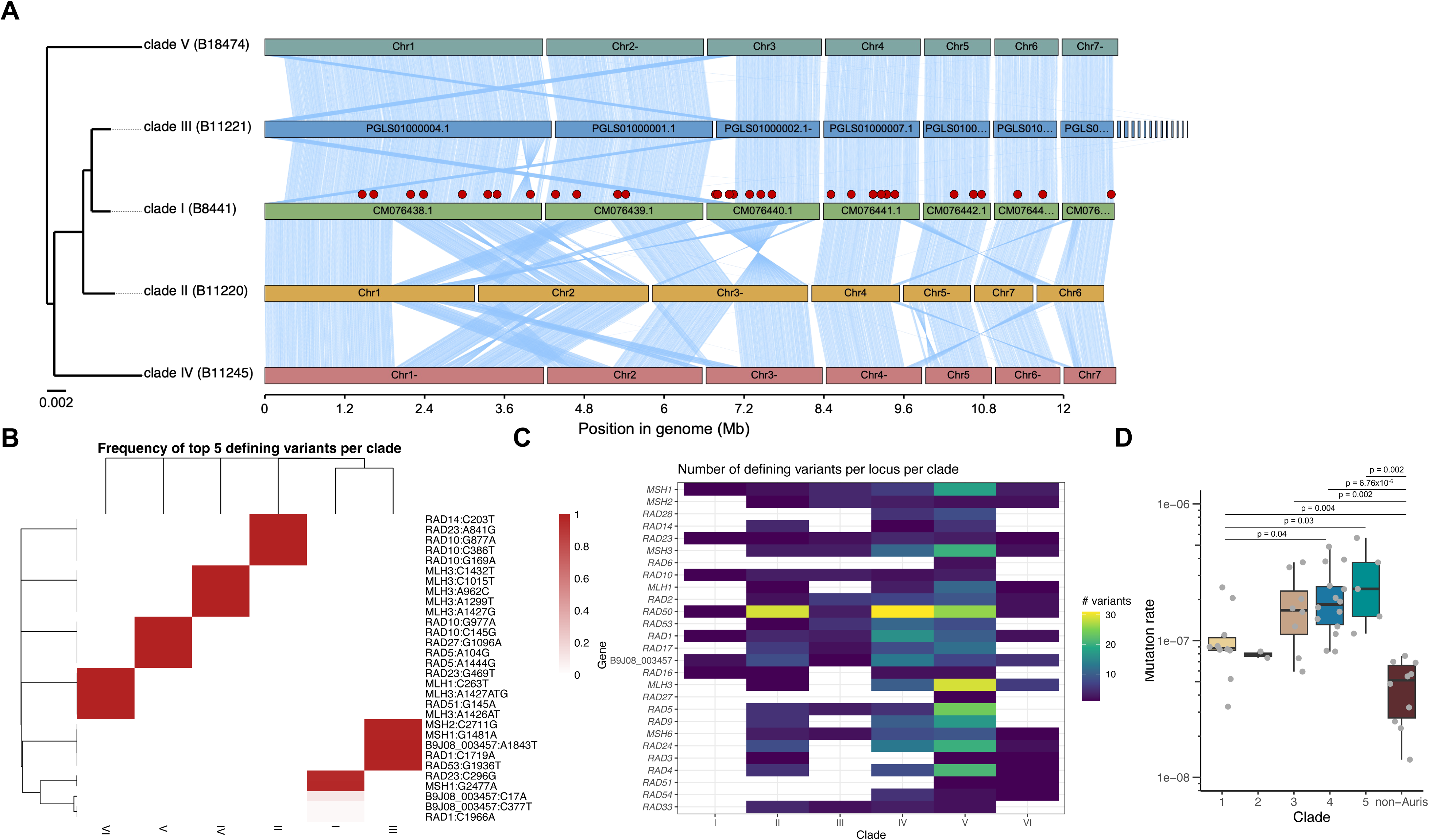
Mutations in MMR genes contribute to elevated mutation rate in *C. auris.* **a)** Synima plot showing reference isolates of five different clades. Red dots represent a MMR gene specifically annotated in clade 1. Only core MMR genes are shown. **b)**Heatmap depicting number of variants in MMR genes per clade **c)** Top five mutations in MMR genes significantly associated with each clade **d)** Luria-Delbrück fluctuation assay suggests *C. auris* has an elevated mutation rate compared to closely related non-*auris Candida* species. Each point shows the mutational frequency of an individual population (*N*D=D30-60). Error bars show SEM and cross bars show median mutational frequency.

Out of 52 core MMR genes, twenty-seven exhibited elevated mutational burdens across all clades, suggesting they function as clade-specific mutational hotspots (Figure 2b). Clade-specific mutational profiles revealed unique missense mutations in MMR genes were significantly associated (Fisher’s exact test and FDR multiple testing correction and odds ratio significant cutoff) with each clade (Figure 2c, Table 2). MMR genes within Clade VI were also significantly associated with an insertion in *MLH3* (A1426AT/G, adj. *P* = 5.84^e-04^). Although some of these genes have been implicated in elevated mutation rates in *C. auris* and other fungal systems^22–24^, the mechanistic contribution of combinations of mutations across MMR genes remains unclear, likely due to epistatic interactions.

**Table 1:** Isolate metadata for initial phenotype screening. Identified MMR genes with missense mutations noted. Minimum Inhibitory Concentrations (MIC) for fluconazole (FCZ), voriconazole (VCZ) and itraconazole (ITZ) provided in mg/mL. N/A = not available.

| Isolate ID | Clade | FCZ MIC | VCZ MIC | ITZ MIC | MMR gene |
| --- | --- | --- | --- | --- | --- |
| CAu_113 | III | >64 | 1 | 0.25 | *MSH1, MSH2, MSH3, MSH6, RAD18, RAD23, RAD53* |
| CAu_117 | III | >64 | 4 | 0.5 | *MSH1, MSH2, MSH3, MSH6, RAD18, RAD23, RAD53* |
| CAu_119 | III | >64 | 4 | 0.5 | *MSH1, MSH2, MSH3, MSH6, RAD18, RAD23, RAD53* |
| CAu_120 | III | >64 | 4 | 0.5 | *MSH1, MSH2, MSH3, MSH6, RAD18, RAD23, RAD53* |
| CAu_121 | III | >64 | 8 | 0.5 | *MSH1, MSH2, MSH3, MSH6, RAD18, RAD23, RAD53* |
| CAu_888 | I | 64 | 0.25 | <0.03 | *MSH1, Mre11, RAD18* |
| CAu_889 | I | 32 | N/A | 0.06 | *MSH1, Mre11, RAD18* |
| CAu_894 | I | 64 | 0.25 | <0.03 | *MSH1, Mre11, RAD18* |
| CAu_895 | I | 32 | 0.5 | <0.03 | *MSH1, Mre11, RAD18* |
| CAu_898 | I | 64 | 0.25 | 0.06 | *MSH1, Mre11, RAD18* |

**Table 2:** Non-synonymous mutations conferring amino acid changes in MMR genes significantly associated (adjusted *p*-value) with each *C. auris* clade.

| Gene | Mutation | Clade I | Clade II | Clade III | Clade IV | Clade V | Clade VI |
| --- | --- | --- | --- | --- | --- | --- | --- |
| *RAD23* | S99C | 2.23<sup>e-</sup><sub>308</sub> |  |  |  |  |  |
|  | I281V |  | 7.49<sup>e-104</sup> |  |  |  |  |
|  | A157S |  |  |  |  |  | 5.75<sup>e-04</sup> |
| *MSH1* | S826N | 2.23<sup>e-</sup><sub>308</sub> |  |  |  |  |  |
|  | R494Q | 2.23<sup>e-</sup><sub>308</sub> |  | 2.23<sup>e-308</sup> |  |  |  |
| *RAD10* | T129M |  | 5.61<sup>e-104</sup> |  |  |  |  |
|  | D57N |  | 5.61<sup>e-104</sup> |  |  |  |  |
|  | D293N |  | 5.61<sup>e-104</sup> |  |  |  |  |
|  | G326D |  |  |  |  | 7.06<sup>e-10</sup> |  |
|  | P49A |  |  |  |  | 7.06<sup>e-10</sup> |  |
| *RAD14* | P68L |  | 8.34<sup>e-104</sup> |  |  |  |  |
| *MSH2* | S904T |  |  | 2.23<sup>e-308</sup> |  |  |  |
| B9J08_003457 | T615S |  |  | 2.23<sup>e-308</sup> |  |  |  |
| *RAD1* | K573N |  |  | 2.23<sup>e-308</sup> |  |  |  |
| *RAD53* | Q646K |  |  | 2.23<sup>e-308</sup> |  |  |  |
| *MLH3* | D478N |  |  |  | 2.23<sup>e-308</sup> |  |  |
|  | D339N |  |  |  | 2.23<sup>e-308</sup> |  |  |
|  | V321G |  |  |  | 2.23<sup>e-308</sup> |  |  |
|  | F433L |  |  |  | 2.23<sup>e-308</sup> |  |  |
|  | L476G |  |  |  | 2.23<sup>e-308</sup> |  |  |
| *RAD27* | P366S |  |  |  |  | 6.05<sup>e-10</sup> |  |
| *RAD5* | H35R |  |  |  |  | 7.49<sup>e-10</sup> |  |
|  | K482E |  |  |  |  | 7.49<sup>e-10</sup> |  |
| *RAD51* | P49S |  |  |  |  |  | 4.65<sup>e-04</sup> |
| *MLH1* | S88N |  |  |  |  |  | 5.47<sup>e-04</sup> |

We assessed the mutation rates across the different clades by performing an adjusted Luria-Delbrück fluctuation assay, which showed *C. auris* has a 4.3-times elevated mutation rate compared to other related *Candida* species (p<0.0001) (Figure 2d). Mutation rates varied markedly among clades: Clades III, IV and V demonstrated the highest rates; Clade IV showed a five-fold increase compared to non-*auris Candida* (p<0.0001), and approximately a two-fold increase relative to Clades I (p=0.04) and II (p=0.02). Mutations were stable as isolates remained 5-FC resistant after passaging in drug-free media (Supplementary Figure 4). This increased mutation rate was consistent using resistance to a clinically used antifungal, 5-flucytosine, and an antifungal currently undergoing clinical trials, manogepix (Supplementary Figure 4).

### Transcriptional rewiring facilitates *C. auris* high salt- and thermo-tolerance

Currently, 21 *C. auris* genome annotations are available on NCBI, of which 12 genomes are of scaffold or chromosome level. Therefore, we annotated an additional 16 genomes across all clades and related non-*C. auris* species. The majority (>87%) of genes were found to be common to all clades (4,865 1:1 core orthologs; between 191 to 249>1:>1 core orthologs; Figure 3a). Pangenome analysis including copy-number variation within orthogroups, showed different genetic composition of *C. auris* isolates to other *Candida* species, as well as clade-specific signatures (Figure 3b, Supp Fig 4). To determine genes that could predict clade, we utilised a Random Forest model (accuracy 96.7%). Within the 100 orthogroups with the lowest mean Decrease Gini, no enrichment of GO or Pfam was found (Supplementary Figure 5). As several genes lacked GO or Pfam domains in the annotation, we used structural similarity (Supplementary Figure 6). This revealed that accessory genes which determined clade, included transporters and cell-wall associated genes.

**Figure 3:**
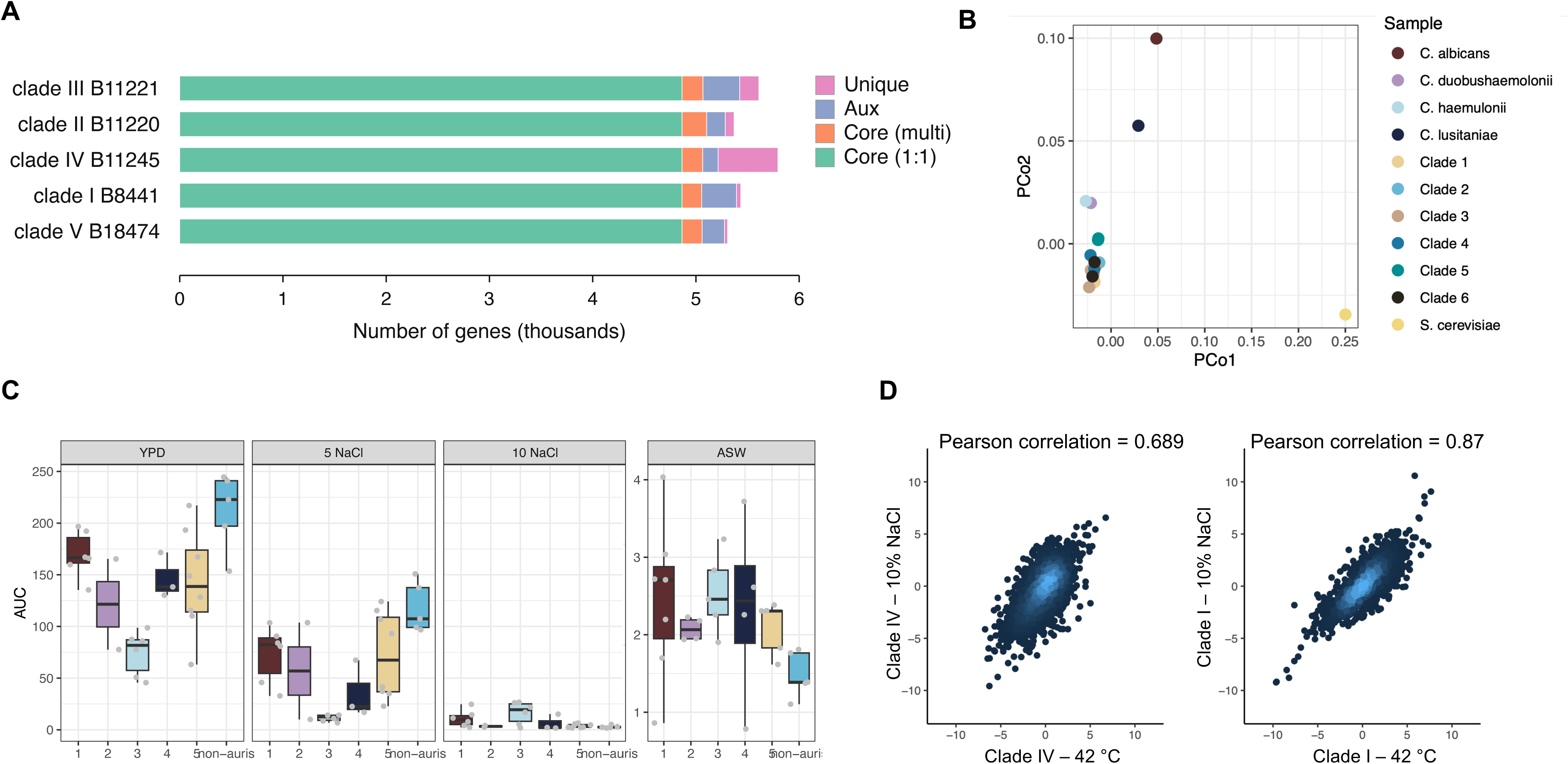
Clade specific adaptation to stress is due to transcriptional rewiring. **a)** Summary of orthologs identified by synteny analysis **b)** PCoA plot for all orthogroups across the different clades and species annotated. **c)** Phenotypic assessment of growth in YPD, under elevated salt (5% and 10% NaCl) and in Artificial Sea Water. **d)** Transcriptional profile of both isolates under elevated salt and temperature conditions. Every dot is one gene; colouring is based on gene density. Pearson correlation for the whole dataset is shown.

As Clade I and III *C. auris* are responsible for the majority of nosocomial infections and outbreaks, we chose to determine if isolates in these clades (*n* = 5 per clade; Table 1) used different growth sources, using Omnilog high-throughput screening carbon and nitrogen sources (PM01 and PM3B, respectively), osmolytes (PM9) and pH (PM10) (Supplementary Figure 7). These isolates all contained missense mutations in MMR genes. Clade III isolates used 39 different carbon sources, while Clade I isolates used only 12 different carbon sources (*p* < 0.01 one-way ANOVA). Differences were also observed between clades specifically under osmotic stress and between pH 3.5 and 10. Therefore, we performed an MTT assay to confirm the relationship between NaCl concentration and growth of Clade I and III *C. auris* isolates. A significant difference (*P* < 0.001) between Clade I and III isolates was observed at 48-hours, with a higher percentage (80%) of Clade III in 5% NaCl solution compared to Clade I (40%). At 10% NaCl, 40% of Clade III survived compared to just 20% Clade I (Supplementary Figure 8).

We performed additional phenotyping to salt tolerance using isolates from 5 clades (*n* = 25) and five non-*auris* isolates (*n*=5) (Figure 3c). In YPD non-*auris* isolates grew better than *C. auris* isolates (P=0.02). Clade III isolates grew slowest in YPD and 5% NaCl compared to other clades (P=0.01). However, at 10% NaCl, Clade III and Clade I showed increased growth compared to other clades and non-*auris* isolates (p=0.035). Clade IV grew better than non-*auris* isolates, although slower than Clade III and I. All clades of *C. auris* were able to grow in artificial seawater (ASW) significantly better than non-*auris* isolates (p=0.04).

As Clade IV tends to cause more invasive disease compared to other *C. auris* clades, we assessed a representative Clade IV isolate (NCPF13048) compared to a representative Clade I isolate (CAu_898, Table 1) using transcriptomic profiling. We sought to investigate the molecular basis of clade-specific differences in salt and temperature tolerance, essential for human infection, through exposure to both high-temperature (42°C) and high-salinity stress (10% w/v NaCl) compared to baseline (37°C in YPD only). Under thermal stress (42°C), the clade IV isolate displayed a markedly different transcriptional response compared to clade I, with 942 genes and 773 genes differentially regulated for clade I and 778 genes and 1128 genes differentially regulated for clade IV under thermal and salt stress, respectively (Supplementary Figure 9).

Exposure to 10% NaCl triggered substantial transcriptional rewiring in both clades, but with pronounced clade-specific patterns (Supplementary Figure 10). The clade IV isolate showed strongly induced genes (>2 log2FoldChange, adjusted p-value<0.05) associated with transcription, translation machinery and tRNA processes, while transmembrane transport, peroxisomes and autophagy were enriched in downregulated genes (<-2 log2FoldChange, adjusted p-value <0.05). Unlike clade IV, upregulated genes in the clade I isolate were enriched for transmembrane transport, while enriched genes such as ribosome biogenesis, translation and RNA binding were shared among both isolates (Supplementary Figure 10). Across both stress conditions, the clade I isolate transcriptome under thermal and salt exposure was more similar (Pearson correlation = 0.87) compared to clade IV isolate (Pearson correlation = 0.689), with a greater number of genes differentially regulated under both conditions and larger fold-change magnitudes compared to Clade IV (Figure 3d). Genes that are upregulated in the clade I isolate under both salt and temperature stress were enriched for transmembrane transport.

Collectively, these results indicate that clade I has undergone extensive transcriptional rewiring that enables more robust adaptation to high salt and high temperature environments. Our transcriptomic data reveal that salt and thermal tolerance in *C. auris* is the result of clade-specific regulatory programmes, with Clade I exhibiting a substantially more flexible and adaptive stress-response architecture than Clade IV. This enhances environmental responsiveness likely contributes to the ecological success of Clade I and may help explain its frequent involvement in clinical outbreaks.

## Discussion

Over the past two decades, *Candidozyma* (*Candida*) *auris* has emerged as a major cause of nosocomial fungal infections worldwide, with repeated outbreaks documented across multiple continents ^1,4,30,31^. To date, six distinct genetic lineages have been identified ^4,32–34^, characterising a unique population structure with high inter-clade diversity and strikingly low intra-clade diversity. *C. auris* exhibits exceptional environmental persistence, stress tolerance, and adhesion³, and is increasingly associated with mutations conferring resistance to all four major antifungal classesD D³ DD. Remarkably, such resistance can arise within individual patients over the course of daysD. Experimental evolution studies have identified the evolution of stable drug resistance in a matter of days in experimentally evolved *C. auris*, with elevated mutation rates proposed as the mechanism for this rapid resistance development ^25^. In this study we demonstrate that *C. auris* now spans all continents, including Antarctica, following its isolation from multiple geographically distinct Antarctic sites. We also demonstrate that mutator phenotypes contribute both to the rapid evolution of antifungal resistance and to the striking global population structure of *C. auris*. Our data suggest that mutator phenotypes confer an evolutionary advantage in certain clades, enhancing adaptability to environmental stressors and potentially influencing host-associated fitness.

The discovery of *C. auris* in Antarctic marine environments raises important questions about its ecological range, dispersal mechanisms, and persistence in low temperature, high salinity niches. The genetic diversity of the Antarctic isolates, combined with evidence of at least two independent introductions, indicates that *C. auris* is not merely a transient contaminant but may survive and persist in these extreme environments, and supports a high-salinity origin niche. While appearing genetically similar to clinical and environmental *C. auris* isolates originating in India and Hong Kong, we cannot currently confirm the transmission routes resulting in *C. auris* deposition in Antarctica. The propensity of *C. auris* to form biofilms on nylon, present in 5% of all manufactured clothing and commonly found in specialist outdoor clothing, suggests human movement could be responsible. However, the recovery of these Antarctic isolates from microplastic debris does not rule out dispersal on ocean currents from the Pacific and Indian oceans, which feed into Antarctic waters through the Antarctic Circumpolar Current (ACC), connecting the world’s oceans.

Our population-scale analysis revealed that across the *C. auris* population, key genes in the mismatch repair (MMR) pathway are significantly more likely to contain mutations, and that mutational profiles are significantly associated with specific genetic clades. Mutations affecting *MSH* family genes, particularly enriched in Clades I and III, are of particular interest, as these proteins recognise base–base mismatches and their disruption is known to substantially elevate mutation rates. Our fluctuation assays support this hypothesis, demonstrating markedly elevated mutation rates in clades bearing MMR-pathway mutations. Although Clade V showed the highest mutation rate across the species, isolates within this clade are rarely recovered; instead, Clades I, III and IV are responsible for most nosocomial outbreaks and infections. Clades I, III and IV were also significantly enriched for mutations in key MMR genes *MSH1*, *MSH2*, and *MLH3,* whereas Clades II and V were significantly enriched for mutations in *RAD* genes. These findings also suggest that combinations of MMR gene mutations may interact epistatically to generate elevated mutation rates, thereby accelerating the accumulation of adaptive mutations, including those conferring antifungal resistance.

Previous studies have reported *C. auris* tolerates high salt concentrations compared to other *Candida* species ^35^. Consistent with this, we observed significantly greater salt tolerance in Clade III compared with Clade I, particularly at 5% and 10% NaCl after 48 hours. Upon expansion of testing to Clades II, IV and V, compared to non-*auris* isolates, we observed Clade IV to grow slower compared to Clades I and III; we therefore used transcriptomic profiling to assess the gene expression changes of fast versus slow growth in salt, using Clade I and Clade IV isolates.

Together, our findings provide new insights into the ecological breadth, evolutionary trajectory, and genetic underpinnings of *C. auris*. The presence of mutator phenotypes across multiple clades, coupled with evidence of environmental persistence and clade specific MMR mutations, suggests that elevated mutation rates have played a central role in the rapid global expansion of this pathogen. Understanding the evolutionary mechanisms shaping *C. auris* diversity will be critical for improving surveillance strategies, anticipating the emergence of resistance, and mitigating the public health threat posed by this rapidly evolving fungus.

## Methods

### Environmental sampling, whole genome sequencing and analysis of *C. auris*

Microplastic particles were collected from seawater using surface sampling approaches designed to minimise contamination. Surface material was obtained by towing a stainless-steel manta net (300–500 µm) across the upper water column and passed through a sequential mesh series (5 mm to 63 µm). All equipment was pre-rinsed, and samples were maintained in glass containers under foil to prevent airborne contamination.

Fungi associated with microplastic surfaces were isolated by resuspending particles in artificial seawater and dislodging biofilms through vortexing. The resulting microbial suspensions were spread onto ChromAgar and plates were incubated until colonies emerged. Individual fungal morphotypes were purified by subculturing onto fresh media. Pure isolates were archived for downstream genomic and phenotypic characterisation.

Genomic DNA was extracted from *C. auris* isolates using a phenol:chloroform protocol optimised for high molecular weight fungal DNA. Briefly, cells were grown overnight in YPD, harvested, and lysed in extraction buffer containing SDS and proteinase K, followed by incubation at 55 °C to ensure complete disruption of the cell wall. Lysates were subjected to sequential phenol:chloroform:isoamyl alcohol extractions to remove proteins and other contaminants, and DNA was precipitated with cold ethanol and sodium acetate before being washed and resuspended in nuclease-free water. RNA was removed by RNase A treatment, and DNA quality was assessed by spectrophotometry and agarose gel electrophoresis to confirm high purity and integrity suitable for whole genome sequencing.

Sequencing libraries were prepared from high quality genomic DNA using standard Illumina protocols, ensuring fragment sizes appropriate for paired-end sequencing. Libraries were indexed and pooled before sequencing on an Illumina platform to generate 150 bp paired end reads, yielding an average depth of coverage sufficient for confident variant discovery across the *C. auris* genome. Raw reads were subjected to initial quality checks using FastQC, and sequencing artefacts including adapter sequences and low-quality bases were removed using Trimmomatic to ensure downstream analyses were performed on high fidelity data. Metadata associated with Antarctic isolates is supplied in Supplementary Table 1.

All short reads were aligned against the *C. auris* GCA_002759435.3 reference assembly using BWA v0.7.17 mem, with further sorting and indexing with SAMtools v1.12. Picard v3.4.0 was used to mark duplicates. GATK v4.4.0.0 was used to call variants and filter single nucleotide polymorphisms (SNPs) and insertions and deletions (INDELs) with the following filters for low-confidence variants: QD < 2.0, FS > 60.0, MQ < 40.0, MQRankSum <-12.5, ReadPosRankSum < −8.0, and SOR > 4.0 for SNPs, and QD < 2.0, FS > 200.0, ReadPosRankSum < −20.0, SOR > 10.0 for INDELs. Variants were annotated with snpEff v5.3a. Depth of coverage was determined using SAMtools and normalised to identify regions of potential aneuploidy.

A further 37 environmental and 91 clinical Clade I isolates, time-matched to the Antarctic isolates, were included for phylogenetic context to assess relationships between these isolates. Phylogenetic trees were first generated with RAxML-NG v1.0.1 with the option to run the maximum likelihood (ML) search and bootstrapping analysis with the GTR+G model. The support option was then run to map the bootstrap values onto the ML trees, which were visualised in FigTree v1.4.5-pre and in R using the ggTree v4.1.1 package.

To assess the temporal signal within these isolates, BEAST v2.7.7 was used. First, PhyloSTemS ^36^ was used to confirm the data was appropriate for tip-dated inference. Root-to-tip regression analysis showed sufficient temporal signal, so bModelTest in BEAST was run to determine the most appropriate GTR+G substitution model with an optimised relaxed clock and Coalescent Bayesian Skyline tree model. BEAST was run for 145,244,000 states with a 10% burn-in and checked for convergence in Tracer v1.7.2. The Maximum Clade Credibility (MCC) tree was generated and visualised in FigTree v1.4.5.

### Alignment of whole-genome sequence reads

All Illumina paired-end whole genome sequencing (WGS) reads for “*Candida auris*” available on NCBI SRA on 23^rd^ August 2024 (*n* = 13,193) were obtained using the ‘Run Selector’ tool ^37^, along with associated metadata. This dataset, along with all project accession numbers, are described in Gifford *et al* ^27^.

A core alignment was created with snippy-core, filtering for a minimum coverage of 10 Mb (nuclear) in batches of 1000 until a successful core phylogeny was identified for each batch. Phylogenetically informative sites were determined using snp-sites v2.5.1 ^38^ across all isolates (*n* = 12,644) with adequate coverage. Approximately maximum likelihood phylogenies were estimated for the nuclear genomes using VeryFastTree v4.0.03 with the GTR rate substitution model ^39^. Midpoint-rooted trees were plotted with ggtree v3.6.2 ^40^ in R v4.2.3.

### Synteny analysis

Ortholog prediction and synteny analysis were performed using Synima v2.0.0 ^28^. Genomes of publicly available *C. auris* isolates with chromosome-level assemblies were obtained from NCBI SRA (clade_III_B11221 (GCA_002775015.1), clade_II_B11220 (GCA_003013715.2), clade_IV_B11245 (GCA_008275145.1), clade_I_B8441 (GCA_002759435.3), clade_V_B18474 (GCA_016809505.1)). Orthologs were inferred using orthomcl v1.4 based on an all-vs-all comparison of peptide sequences computed with diamond v2.1.6 ^41^ using the parameters max_target_seqs=250, evalue=1e-10, diamond_sensitivity=fast.

Orthogroups assigned by Synima were classified into core, accessory, and unique categories. 4865 single-copy core orthologs were identified and used to construct a phylogenetic tree. Each orthogroup of single-copy orthologs was aligned separately using MUSCLE v5.3.osx64 ^42^ with default settings. All alignments were concatenated into a single FASTA, and an ‘approximately maximum-likelihood’ tree was inferred using FastTree v2.1.11 SSE3 ^43^.

Synteny blocks were identified as chains of ≥ 4 orthologous genes using DAGChainer ^44^, and visualised using Synima.

### Orthogroup analysis

Annotations, if available, were also obtained, or generated using FunAnnotate v1.8.14. Orthogroups were determined using OrthoFinder v3.0. A Random Forest model to determine orthogroups with the best predictive power of clade (lowest Gini score) was generated in R v4.5.2 using the RandomForest package v4.7-1.2. The top 100 orthogroups with the lowest Gini score were visualised using pheatmap in R.

### Biofilm formation on plastic substrates

Following biofilm formation for 24 hours in YPD+Salt on sterile, weight-matched plastic substrates, culture supernatants and non-adherent cells were carefully aspirated. Biofilms were washed and air-dried, prior to staining with 200 µL of 0.1% (w/v) crystal violet solution dHDO for 30 min. Excess stain was removed by aspiration, and biofilms washed.

Bound crystal violet was solubilised with 10% (v/v) acetic acid prepared in dHDO for 10 min at room temperature with gentle agitation. The resulting suspension was transferred to a fresh 96-well plate and total biofilm biomass was quantified by measuring absorbance at 595 nm (ODDDD) using a microplate reader.

### Phenotyping

Initial screening was performed on ten isolates (Table 1). Isolates were revived *via* culturing on Sabouraud dextrose agar plates (ThermoFisher, USA) and incubated at 37°C for 24-48 hours until characteristic white, smooth colonies were observed. Individual colonies were picked into FF Inoculating Fluid (Biolog, Hayward, CA, USA) to achieve a required density of 62-65%. High-throughput phenotype screening was performed by incubating plates in the Omnilog system (Biolog, Hayward, CA, USA) with PM01 (carbon source), PM3B (nitrogen source), PM9 (osmolytes) and PM10 (pH) microplates (Biolog, Hayward, CA, USA). All components were well mixed, and 200 µL was pipetted into each well in the microplate.

Microplates were incubated in the Biolog Omnilog for 120 hours at 37°C, with readings every 15 minutes. Growth data was recorded using the Biolog Data Analysis software v1.7 (Biolog, Hayward, CA, USA). Data was analysed in RStudio v4.3.2 using package ‘opm’ and visualised using ‘Metaboanalyst’. Raw data was normalised, and a *p*-value threshold set to 0.05 with unequal group variation.

To follow up the growth in sodium chloride (NaCl) conditions, an MTT assay was performed. 8 g NaCl was dissolved in 40 mL RPMI 1640 liquid to obtain 20% NaCl solution, and filtered using Nalgene filtration (Thermos Fisher Scientific, USA) and added to a 96-well plate for eight technical duplicates per condition. Plates were incubated at 37°C for up to 48 hours. 5 mg/mL MTT 25 µL per well was added with further incubation for 3 hours at 37°C. All media was discarded prior to adding 200 µL 5% hydrochloric acid (Atom Scientific, Manchester, UK) and 2-Propanol (Sigma-Aldrich, USA) solution per well to end the MTT assay. Plates were gently agitated on an orbital shaker for 30 mins at room temperature. The solution was mixed *via* pipetting until fully dissolved. Read plate absorbance at optical density 570 nm via microplate absorbance read (Bio-RAD, Berkeley, CA, USA) within one hour.

Growth curve analysis was performed by reviving isolates on Sabouraud dextrose agar plates and incubated at 37°C for 24-48 hours until smooth colonies were observed. Individual isolates were grown in liquid YPD at 150 rpm at 37°C until OD600=0.5 was achieved. Artificial seawater medium was created as described ^45^. 200 µL of medium (YPD, YPD + indicated percentage of NaCl, or other indicated medium and temperatures) was inoculated in a 96-well plate (Starlab, UK) and grown at 37°C on a Tecan Sunrise plate reader. Area Under the Curve was determined in R v4.5.2.

### Fluctuation assays

Unless otherwise stated, all isolates were cultured on Sabouraud dextrose (SAB) agar (Formedium, UK) at 37°C. 5-fluorocytosine (5-FC) powder was obtained from Fluorochem. Manogepix (E1210) was obtained from Selleckchem. Each drug was initially dissolved in DMSO (Sigma-Aldrich, USA) at 1000-times the highest testing concentration. Specifically, the stock solution of 5-FC was prepared at a concentration of 100 mg/mL, and the stock solution of manogepix was prepared at a concentration of 0.78 mg/mL. The prepared sterile drug stock solution was carefully sealed and stored in a −20°C freezer until required.

Isolates were streaked onto SAB solid medium and incubated at 37°C for 24-48 hours. After incubation, a single colony was picked from the plates using a sterile inoculating loop into 50 mL Falcon tubes (Sarstedt, UK) containing 25 mL SAB liquid medium. 100 µL of this was inoculated in each of 60 wells of a 96-well plate (Starlab, UK). Cultures were incubated at 130 rpm at 37°C for 16 hours. Cultures were inoculated into each well of a six-well plate containing antifungals and SAB agar medium: 1) SAB medium plus 100 mg/L 5-FC and 2) SAB medium plus 0.78 mg/L manogepix. Culture plates were sealed with parafilm and incubated at 37°C for 48 hours following counting of resistant colonies.

The number of mutation events (*m*) was estimated using the maximum likelihood methods of the Luria-Delbrück distribution implemented in the ‘flan’ R package v1.0. The total number of cells (*Nt*) was measured by performing serial dilution on two wells to count total colonies on SAB agar. The mutation rate for each isolate was calculated by dividing *m* by the final population size *Nt,* resulting in a mutation rate of *m/Nt*.

For E-test and disc assay validation, a single colony was picked and inoculated into a 50 mL Falcon tube containing 5 mL SAB liquid medium and incubated at 37°C 130 rpm for 16 hours. 50 µL of culture was transferred to a new 50 mL Falcon tube and diluted 100-times by adding 5 mL of SAB liquid medium. Using a sterile cotton swab, the prepared *C. auris* suspension was evenly spread on the surface of a SAB agar plate, ensuring complete coverage of the plate. A 5-FC E-test strip (Liofilchem, Italy) was placed on the agar surface using sterile forceps. For the disc assay, after evenly spreading on a SAB agar plate, a disc (GE Healthcare, UK) was positioned with sterile forceps. 10 µL of 780 mg/L manogepix was applied to the disc. The inoculated plates with either the E-test strip or disc were incubated inverted at 37°C for 24-48 hours.

### RNA sequencing

Isolates were revived *via* culturing on SAB agar plates (ThermoFisher, USA) and incubated at 37°C for 24-48 hours until characteristic white, smooth colonies were observed. A single colony was picked and inoculated into a 1 mL Eppendorf tube containing either YPD or YPD + 10% NaCl. The culture was incubated at 37°C or 42°C with shaking at 130 rpm for 16 hours. RNA was extracted using the YeaStar RNA extraction kit (Zymo, UK). RNA sequencing was performed by Novogene, UK on a NovaSeq X (2×150bp).

Raw fastq files were obtained and quality control was performed using FastQC v0.12.1, followed by adapter trimming using Trimmomatic v0.38. Reads were aligned to the *C. auris* clade 1 reference genome (B8441) or clade 4 reference genome (B11245). Differential expression analysis was performed using DESeq2 v1.18.1. Differentially expressed genes were considered (<-2 or >2 and adjusted p-value < 0.05) Enrichment analyses were performed using GO terms in FungiFun v3.

### Statistics

All statistics were performed in R v4.2.3. We assessed whether the presence of a mutation within an MMR gene is non-randomly associated with membership in a *C. auris* clade using Fisher’s exact test and FDR multiple testing correction and odds ratio significant cutoff.

## References

1. Satoh, K., et al. *Candida auris* sp. nov., a novel ascomycetous yeast isolated from the external ear canal of an inpatient in a Japanese hospital. Microbiol. Immunol. 53, 41–44 (2009).

2. Jesudason, T. UKHSA publishes updated guidance for C auris. Lancet Microbe 6, 101151 (2025).

3. Gifford, H., Rhodes, J. & Farrer, R. A. The diverse genomes of *Candida auris*. Lancet Microbe 100903 (2024) doi:10.1016/S2666-5247(24)00135-6.

4. Lockhart, S. R. et al. Simultaneous emergence of multidrug resistant *Candida auris* on three continents confirmed by whole genome sequencing and epidemiological analyses. Clin. Infect. Dis. 64, 134–140 (2017).

5. Muñoz, J. F. et al. Clade-specific chromosomal rearrangements and loss of subtelomeric adhesins in *Candida auris*. Genetics 218, iyab029 (2021).

6. Chowdhary, A., Sharma, C. & Meis, J. F. *Candida auris*: A rapidly emerging cause of hospital-acquired multidrug-resistant fungal infections globally. PLoS Pathog. 13, e1006290 (2017).

7. Jacobs, S. E., et al. *Candida auris* Pan-Drug-Resistant to Four Classes of Antifungal Agents. Antimicrob. Agents Chemother. e00053–22 (2022) doi:10.1128/aac.00053-22.

8. Wang, Y. & Xu, J. Population genomic analyses reveal evidence for limited recombination in the superbug *Candida auris* in nature. Comput. Struct. Biotechnol. J. 20, 3030–3040 (2022).

9. Sharma, M. & Chakrabarti, A. On the Origin of *Candida auris*: Ancestor, Environmental Stresses, and Antiseptics. mBio 11, e02102–20 (2020).

10. Casadevall, A., Kontoyiannis, D. P. & Robert, V. Environmental *Candida auris* and the Global Warming Emergence Hypothesis. mBio 12, e00360–21 (2021).

11. Casadevall, A., Kontoyiannis, D. P. & Robert, V. On the Emergence of *Candida auris*: Climate Change, Azoles, Swamps, and Birds. mBio 10, 41–7 (2019).

12. Lima, S. L. et al. Increasing Prevalence of Multidrug-Resistant *Candida haemulonii* Species Complex among All Yeast Cultures Collected by a Reference Laboratory over the Past 11 Years. J. Fungi 6, 110 (2020).

13. Akinbobola, A., Kean, R. & Quilliam, R. S. Plastic pollution as a novel reservoir for the environmental survival of the drug resistant fungal pathogen *Candida auris*. Mar. Pollut. Bull. 198, 115841 (2024).

14. Welsh, R. M. et al. Survival, Persistence, and Isolation of the Emerging Multidrug-Resistant Pathogenic Yeast *Candida auris* on a Plastic Health Care Surface. J. Clin. Microbiol. 55, 2996–3005 (2017).

15. Arora, P. et al. Environmental Isolation of *Candida auris* from the Coastal Wetlands of Andaman Islands, India. mBio 12, e03181–20 (2021).

16. Chow, N. A. et al. Tracing the Evolutionary History and Global Expansion of *Candida auris* Using Population Genomic Analyses. mBio 11, e03364–19 (2020).

17. Naicker, S. D. et al. Clade distribution of *Candida auris* in South Africa using whole genome sequencing of clinical and environmental isolates. Emerg. Microbes Infect. 10, 1300–1308 (2021).

18. Maphanga, T. G., et al. *In Vitro* Antifungal Resistance of *Candida auris* Isolates from Bloodstream Infections, South Africa. Antimicrob. Agents Chemother. 65, e00517–21 (2021).

19. Yadav, A., et al. *Candida auris* on Apples: Diversity and Clinical Significance. mBio 13, e00518–22 (2022).

20. Massic, L. et al. Detection of five instances of dual-clade infections of *Candida auris* with opposite mating types in southern Nevada, USA. Lancet Infect. Dis. 23, e328–e329 (2023).

21. Lin, Z., Nei, M. & Ma, H. The origins and early evolution of DNA mismatch repair genes—multiple horizontal gene transfers and co-evolution. Nucleic Acids Res. 35, 7591–7603 (2007).

22. Rhodes, J. et al. A Population Genomics Approach to Assessing the Genetic Basis of Within-Host Microevolution Underlying Recurrent *Cryptococcal Meningitis* Infection. G3 Genes Genomes Genet. 7, 1165–1176 (2017).

23. Billmyre, R. B., Clancey, S. A. & Heitman, J. Natural mismatch repair mutations mediate phenotypic diversity and drug resistance in *Cryptococcus deuterogattii*. 1–52 (2017) doi:10.1101/141507.

24. Billmyre, R. B., Clancey, S. A., Li, L. X., Doering, T. L. & Heitman, J. Hypermutation in *Cryptococcus* reveals a novel pathway to 5-fluorocytosine (5FC) resistance. 15, 45–34 (2019).

25. Burrack, L. S., Todd, R. T., Soisangwan, N., Wiederhold, N. P. & Selmecki, A. Genomic Diversity across *Candida auris* Clinical Isolates Shapes Rapid Development of Antifungal Resistance *In Vitro* and *In Vivo*. mBio 13, e00842–22 (2022).

26. Bottery, M. J. et al. Elevated mutation rates in multi-azole resistant *Aspergillus fumigatus* drive rapid evolution of antifungal resistance. Nat. Commun. 15, 10654 (2024).

27. Gifford, H. et al. Global genomic epidemiology of *Candida auris*: analysis of 12,644 whole genome sequences from 1997-2024. bioRxiv (2026).

28. Farrer, R. A. Synima: a Synteny imaging tool for annotated genome assemblies. BMC Bioinformatics 18, 507 (2017).

29. Muñoz, J. F. et al. Genomic insights into multidrug-resistance, mating and virulence in *Candida auris* and related emerging species. Nat. Commun. 9, 5346 (2018).

30. Schelenz, S. et al. First hospital outbreak of the globally emerging *Candida auris* in a European hospital. Antimicrob. Resist. Infect. Control 1–7 (2016) doi:10.1186/s13756-016-0132-5.

31. Zhu, Y. Laboratory Analysis of an Outbreak of *Candida auris* in New York from 2016 to 2018: Impact and Lessons Learned. J. Clin. Microbiol. 58, e02083–19 (2020).

32. Spruijtenburg, B. et al. Confirmation of fifth *Candida auris* clade by whole genome sequencing. Emerg. Microbes Infect. 11, 2405–2411 (2022).

33. Khan, T. et al. Emergence of the novel sixth *Candida auris* Clade VI in Bangladesh. Microbiol. Spectr. 12, e03540–23 (2024).

34. Suphavilai, C. et al. Detection and characterisation of a sixth *Candida auris* clade in Singapore: a genomic and phenotypic study. Lancet Microbe 100878 (2024) doi:10.1016/S2666-5247(24)00101-0.

35. Brandt, P. et al. High-Throughput Profiling of *Candida auris* Isolates Reveals Clade-Specific Metabolic Differences. Microbiol. Spectr. 11, e00498–23 (2023).

36. Doizy, A., Prin, A., Cornu, G., Chiroleu, F. & Rieux, A. *PhyloSTemS:* A New Graphical Tool to Investigate Temporal Signal of Heterochronous Sequences at Various Evolutionary Scales. doi:10.22541/au.159050344.45788558.

37. Wheeler, D. L. et al. Database resources of the National Center for Biotechnology Information. Nucleic Acids Res. 36, D13–D21 (2007).

38. Page, A. J. et al. SNP-sites: rapid efficient extraction of SNPs from multi-FASTA alignments.

39. Piñeiro, C., Abuín, J. M. & Pichel, J. C. Very Fast Tree: speeding up the estimation of phylogenies for large alignments through parallelization and vectorization strategies. Bioinformatics 36, 4658–4659 (2020).

40. Yu, G., Smith, D. K., Zhu, H., Guan, Y. & Lam, T. T. GGTREED: an R package for visualization and annotation of phylogenetic trees with their covariates and other associated data. Methods Ecol. Evol. 8, 28–36 (2017).

41. Buchfink, B., Xie, C. & Huson, D. H. Fast and sensitive protein alignment using DIAMOND. Nat. Methods 12, 59–60 (2015).

42. Edgar, R. C. MUSCLE: multiple sequence alignment with high accuracy and high throughput. Nucleic Acids Res. 32, 1792–1797 (2004).

43. Price, M. N., Dehal, P. S. & Arkin, A. P. FastTree: Computing Large Minimum Evolution Trees with Profiles instead of a Distance Matrix. Mol. Biol. Evol. 26, 1641–1650 (2009).

44. Haas, B. J., Delcher, A. L., Wortman, J. R. & Salzberg, S. L. DAGchainer: a tool for mining segmental genome duplications and synteny. Bioinformatics 20, 3643–3646 (2004).

45. Smith, C. & Ferrer-Gonzalez, F. Artificial Seawater Medium protocol. (2018) doi:10.17504/protocols.io.jvccn2w.

