## Supplementary figures and images for "Antarctic marine microplastics reveals environmental persistence and rapid evolution of *Candida auris*"

### Supplementary Figure 1

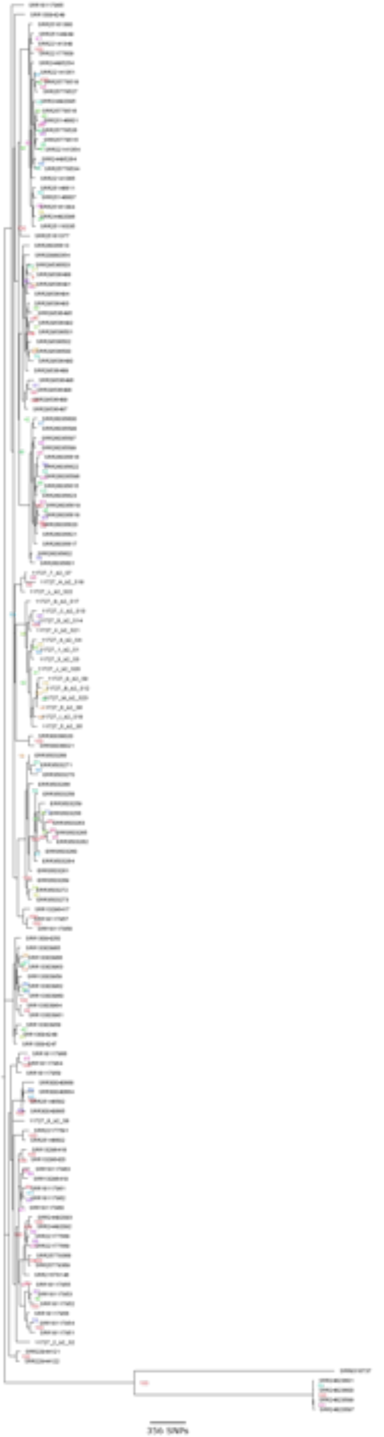

### Supplementary Figure 2

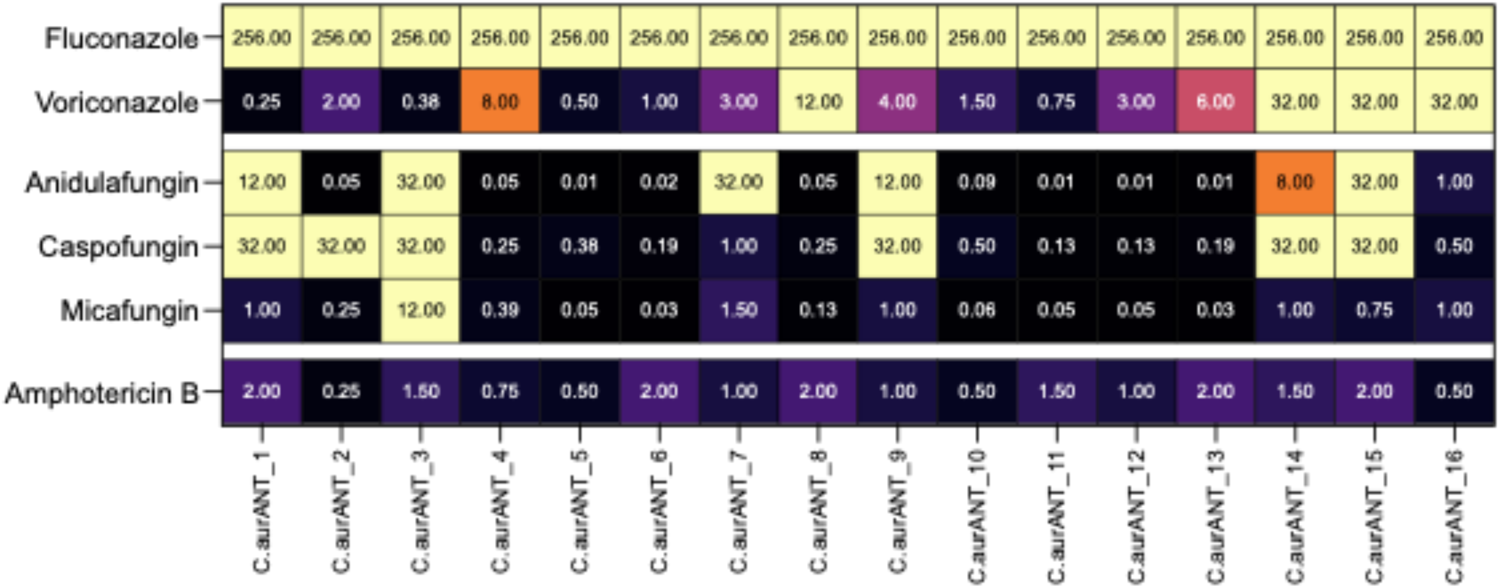

### Supplementary Figure 3

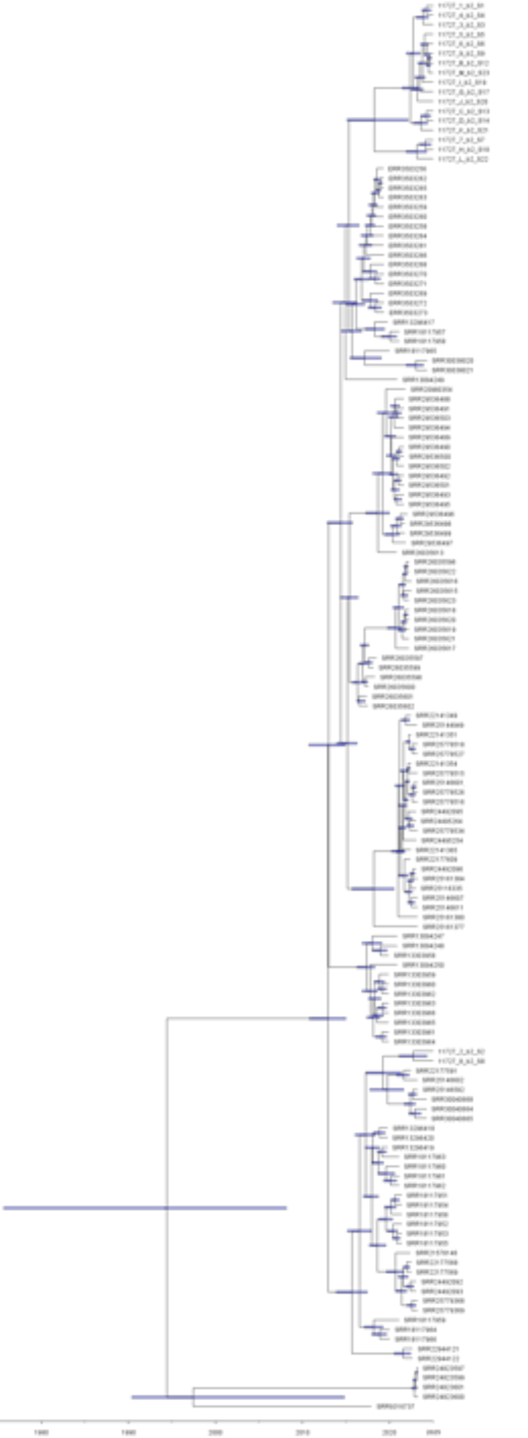

### Supplementary Figure 4

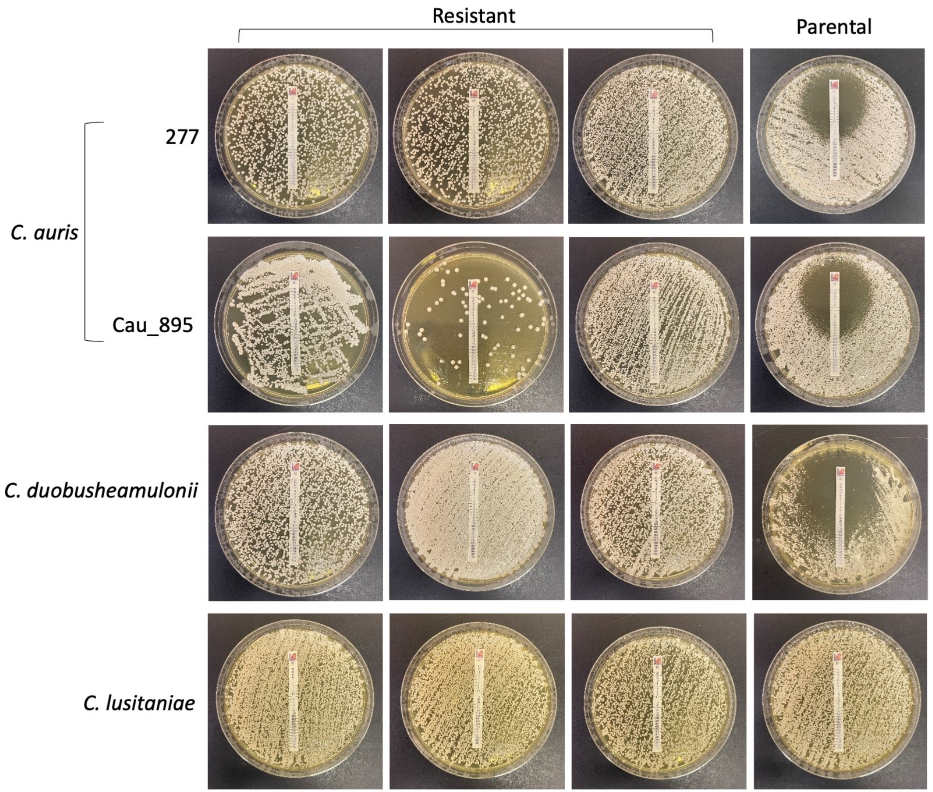

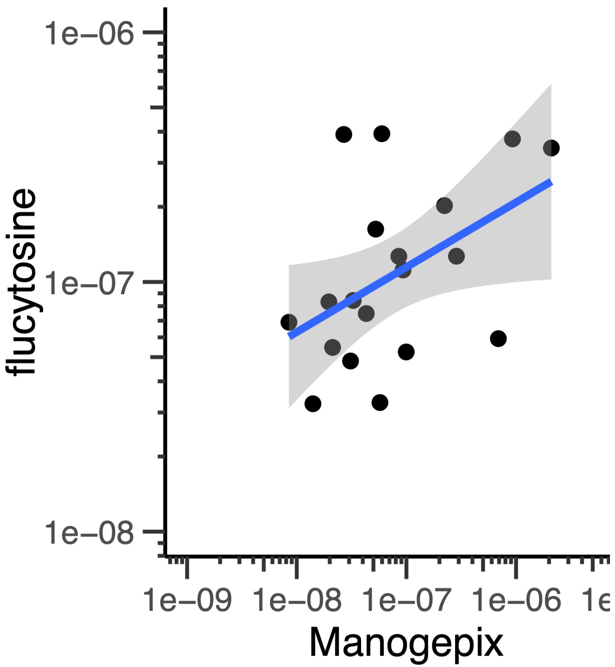

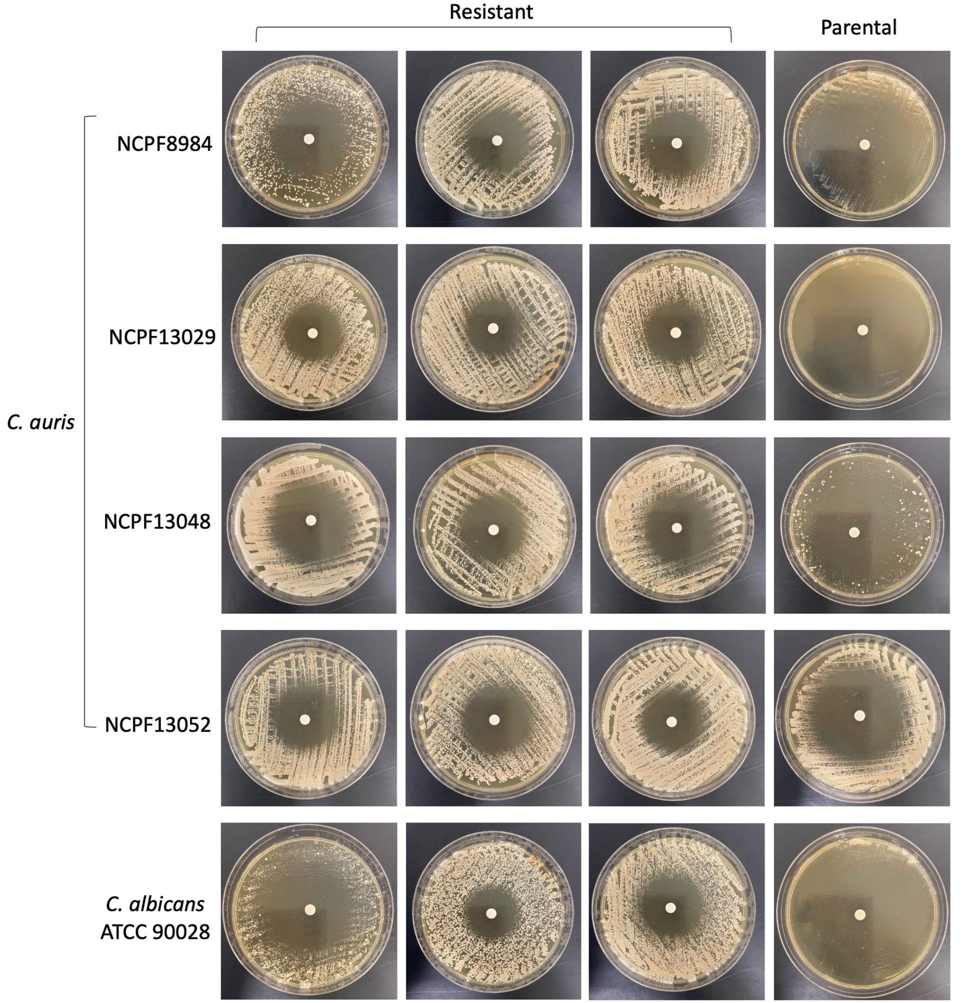

### Supplementary Figure 5

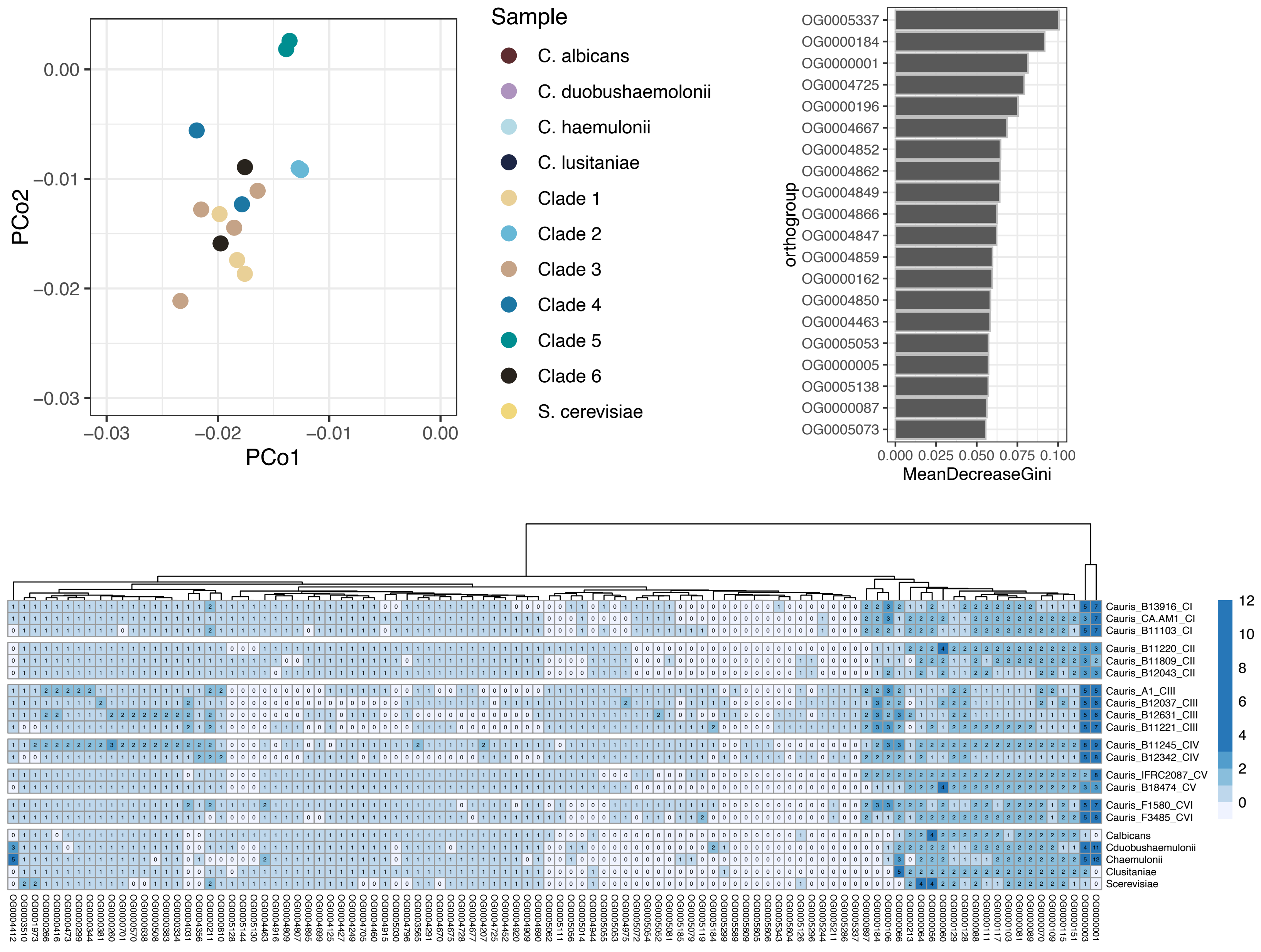

### Supplementary Figure 6

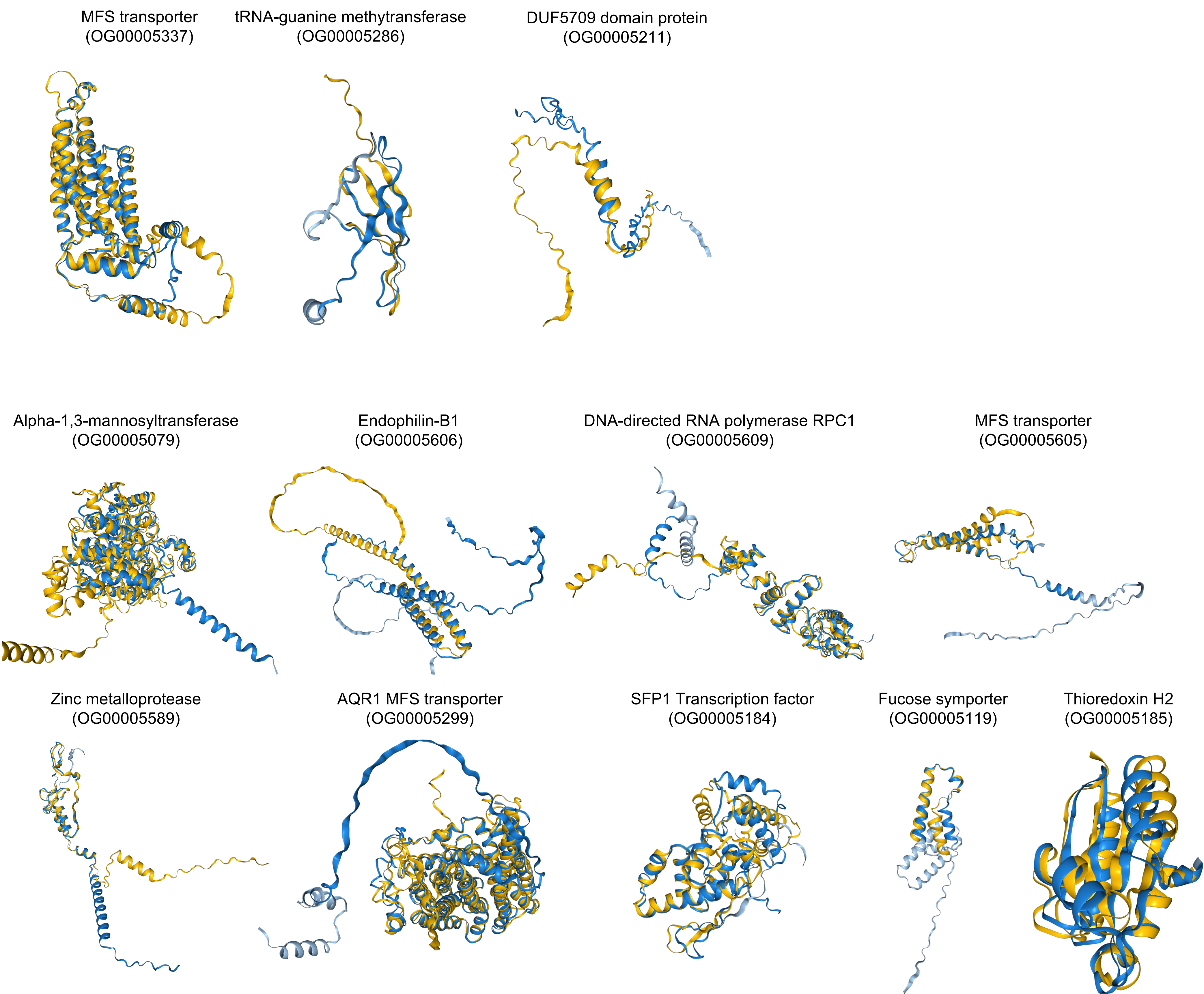

### Supplementary Figure 7

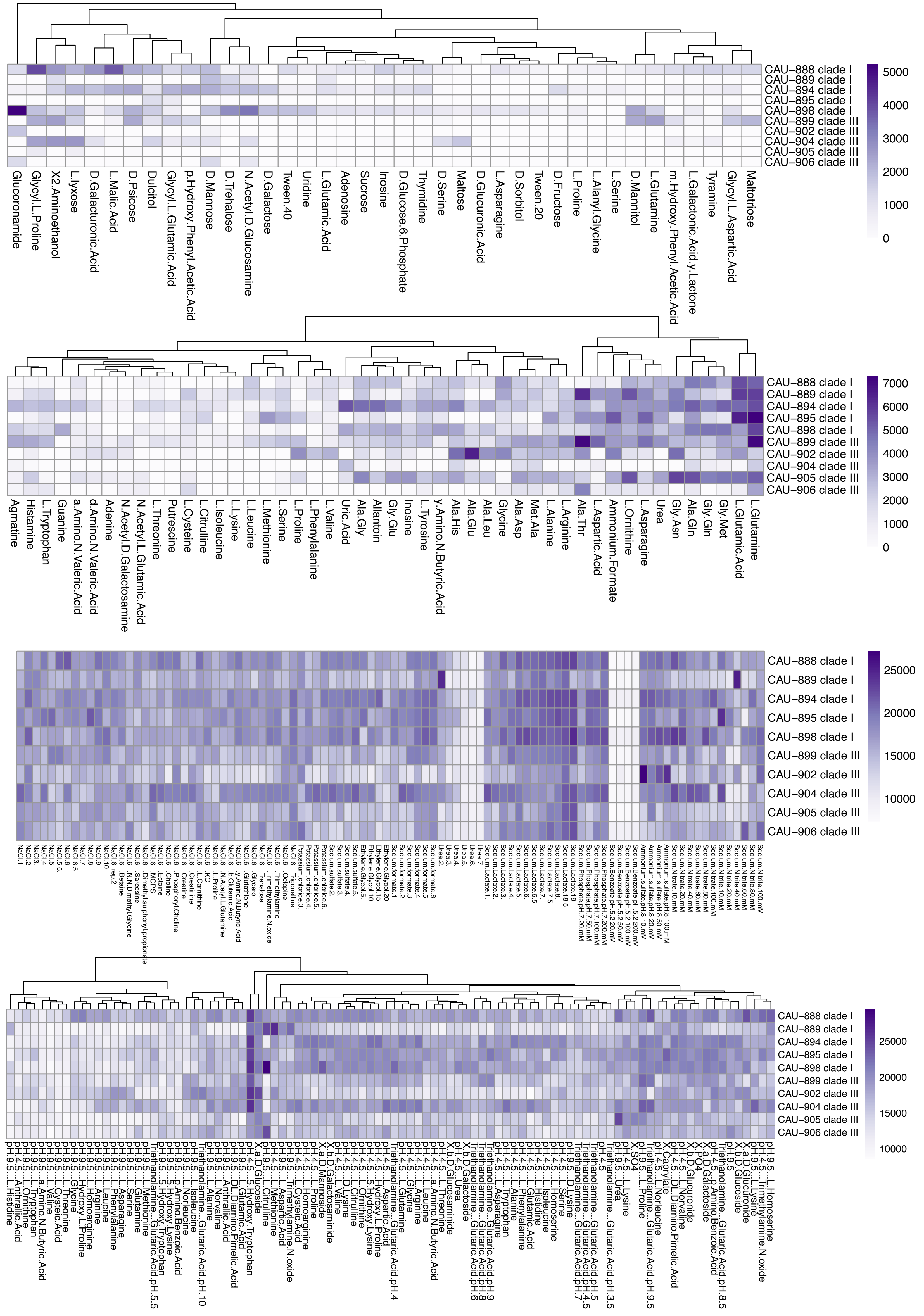

### Supplementary Figure 8

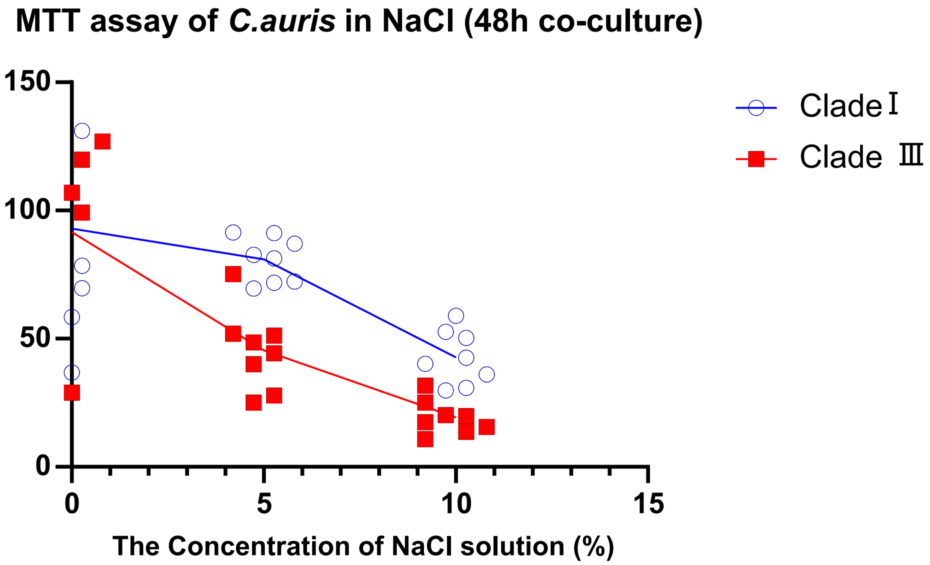

### Supplementary Figure 9

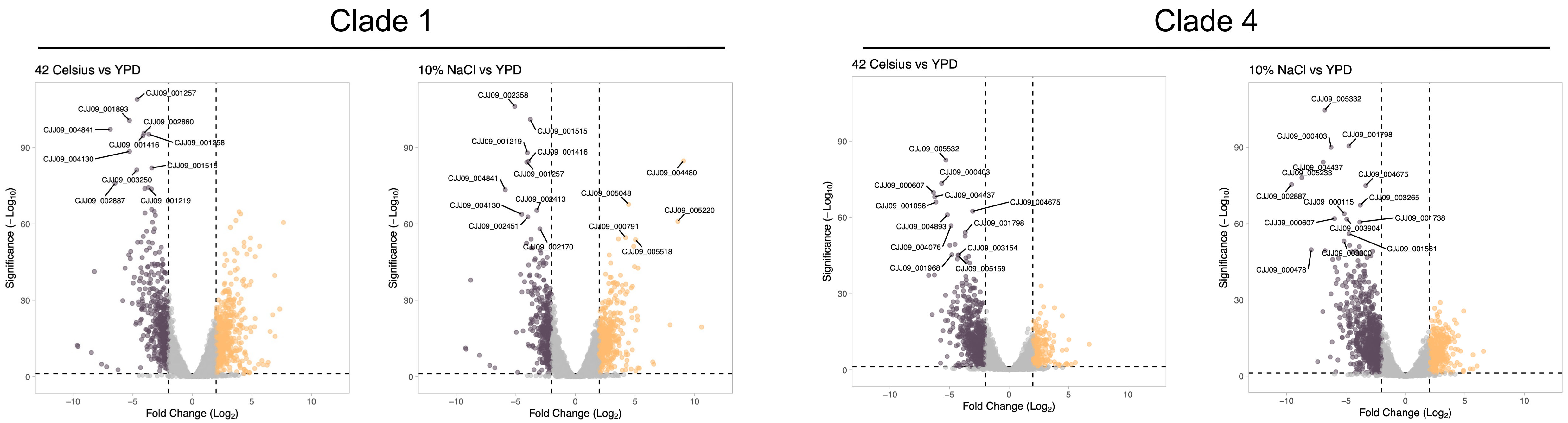

### Supplementary Figure 10

#### Clade IV 10% NaCl

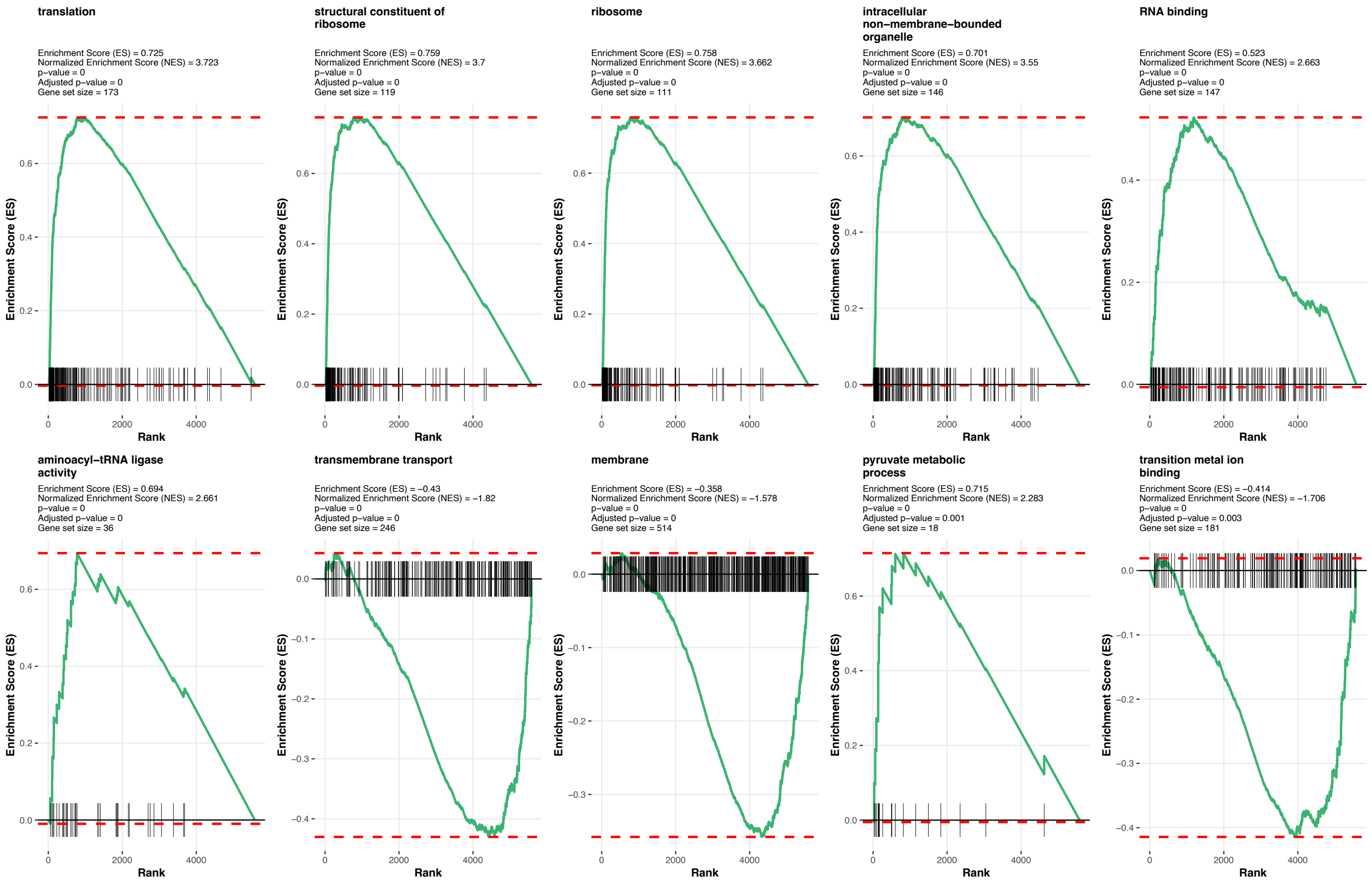


#### Clade I 10% NaCl

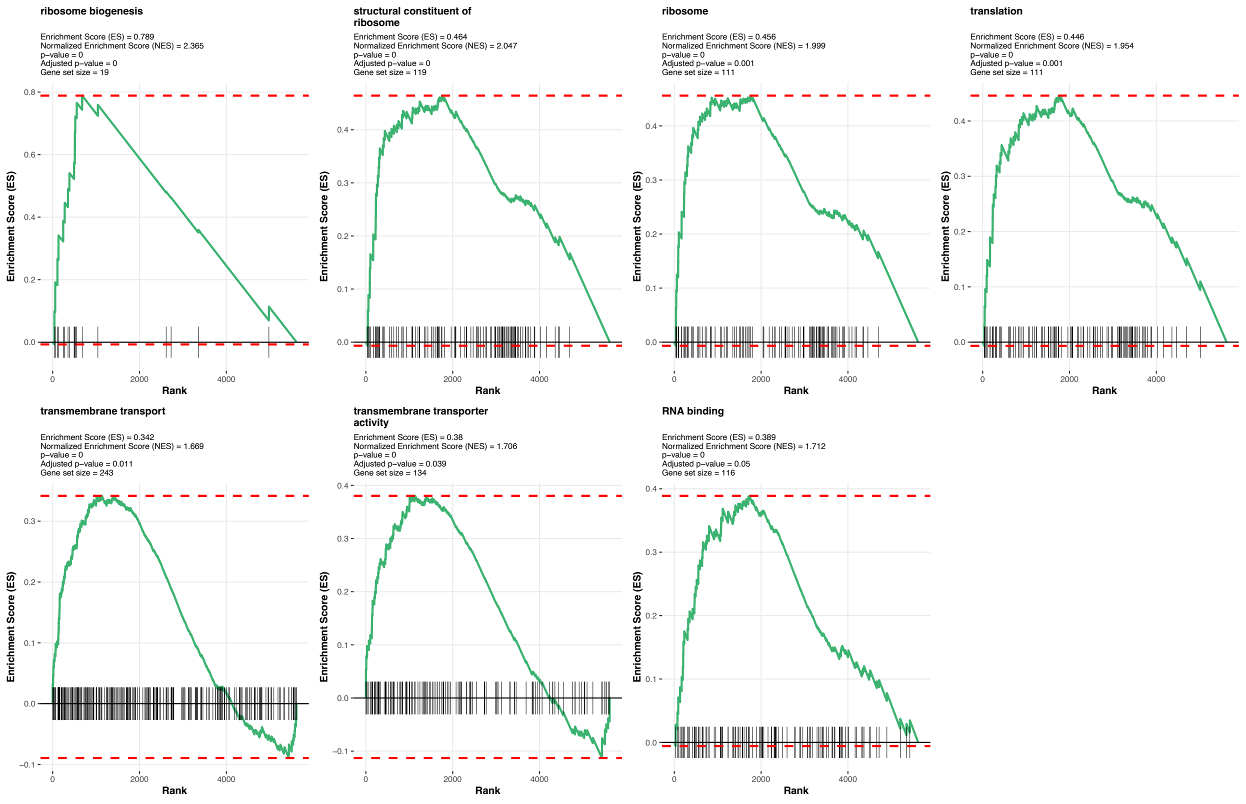
