## Supplementary Information for "Antarctic marine microplastics reveals environmental persistence and rapid evolution of *Candida auris*"

Supplementary Table 1: Sampling locations and statistics from whole genome alignment of 19 isolates from Antarctica against the *C. auris* B8441 v3 reference genome, and identified known drug resistance mutations.

| Isolate ID | Sampling location | Coverage (*x*) | % genome covered | *ERG11* | *FKS1* | *FLO8* |
| --- | --- | --- | --- | --- | --- | --- |
| **C.aurANT_19** | Isolate not banked | 180 | 99.2 | Y132F |  |  |
| **C.aurANT_17** | Isolate not banked | 138 | 99.3 | K143R |  | Q142* |
| **C.aurANT_18** | Isolate not banked | 133 | 99.2 | Y132F | F635L |  |
| **C.aurANT_1** | Paulet Island | 122 | 99.2 | Y132F |  |  |
| **C.aurANT_2** | Paulet Island | 127 | 99.2 | Y132F |  |  |
| **C.aurANT_3** | Paulet Island | 147 | 99.2 | Y132F |  |  |
| **C.aurANT_4** | Deception Island | 152 | 99.1 | Y132F |  |  |
| **C.aurANT_5** | Deception Island | 164 | 99.2 | K143R |  |  |
| **C.aurANT_6** | Deception Island | 145 | 99.2 | Y132F | F635Y |  |
| **C.aurANT_7** | Deception Island | 130 | 99.2 | Y132F |  |  |
| **C.aurANT_8** | Spert Island | 119 | 99.3 | Y132F |  |  |
| **C.aurANT_9** | Spert Island | 131 | 99.3 | Y132F |  |  |
| **C.aurANT_10** | Palmer Station | 127 | 99.2 | Y132F |  |  |
| **C.aurANT_11** | Palmer Station | 119 | 99.2 | Y132F |  |  |
| **C.aurANT_12** | Palmer Station | 114 | 99.2 | Y132F |  |  |
| **C.aurANT_13** | Palmer Station | 108 | 99.2 | Y132F |  |  |
| **C.aurANT_14** | Port Lockroy | 122 | 99.3 | Y132F |  |  |
| **C.aurANT_15** | Port Lockroy | 100 | 99.3 | Y132F |  |  |
| **C.aurANT_16** | Port Lockroy | 142 | 99.2 | Y132F |  |  |

**Supplementary Figure 1:** Maximum likelihood phylogeny of Antarctic isolates plus additional clinical and environmental Clade I *C. auris* isolates for contextualisation, with bootstrap support. Branch length represents the average number of SNPs, and branch labels indicate the bootstrap support.

**Supplementary Figure 2:** Minimum inhibitory concentration for Antarctic *C. auris* isolates for azoles (fluconazole and voriconazole), echinocandins (micafungin, caspofungin and anidulafungin) and amphotericin B.

**Supplementary Figure 3:** Maximum clade credibility phylogeny of Antarctic plus additional clinical and environmental Clade I *C. auris* isolates, with posterior distribution shown.

**Supplementary Figure 4: Mutation rate varies across clades.** a) Validation of stable resistance to 5-FC after passaging in drug free media and susceptibility testing using E-test. B) Mutation rates correlated across using resistance to 5-FC and manogepix as a marker. C) Manogepix resistance was verified to be stable after passaging in drug-free media followed by susceptibility determination.

**Supplementary Figure 5:** **Orthogroup analysis of different clades. a)** Zoomed PcoA on only *Candida auris* isolates showed that orthogroup of isolates can separate clades. **b)** Top 20 orthogroups as identified by highest Decrease in Gini score from the Random Forest on orthogroups to determine clade in *C. auris*. **c)** Heatmap of top 100 orthogroups to determine clade as defined by the Random Forest.

**Supplementary Figure 6: Foldseek analysis of clade defining genes**. Alphafold3 was used on proteins that defined clade, shown in yellow. Foldseek was used to find structural similar proteins, which are shown in blue.

**Supplementary Figure 7: Omnilog analysis of five clade I and five clade III isolates.** AUCs are plotted for Omnilog PM1, PM3 (for both these normalised to negative control) and for PM9 and PM10 (AUC from each growth curve).

**Supplementary Figure 8: MTT cell variability of *C. auris* co-cultured with sodium chloride (NaCl) solution**. The red line represents clade Ⅲ and the blue line represents clade . Each clade was assessed using eight technological replicates in each condition. Four biological replicates were assessed in 24-hour co-culture environment.

**Supplementary Figure 9:** **RNA-seq under elevated salt and temperature.** Volcano plots of a clade I and clade IV isolate under elevated salt and temperature conditions normalised to YPD. Differential expression was assessed using DESeq2 and cut-offs of -2 and 2 (both adjusted p-value<0.05) were considered significant.

**Supplementary Figure 10: Enrichment analysis of transcriptome under elevated salt.** Enrichment analysis was performed using GSEA on FungiFun3 with the whole transcriptome as input data. Top 10 categories are shown here.
